# E2F3-dependent activation of FAM111B restricts mouse cytomegalovirus replication in primate cells

**DOI:** 10.1101/2024.08.02.606359

**Authors:** Eleonore Ostermann, Laura-Marie Luoto, Michaela Clausen, Sanamjeet Virdi, Wolfram Brune

## Abstract

Cytomegaloviruses are highly species-specific as they replicate only in cells of their own or a closely related species. For instance, human cytomegalovirus cannot replicate in rodent cells, and mouse cytomegalovirus (MCMV) cannot replicate in human and monkey cells. However, the mechanisms underlying the host species restriction remain poorly understood. We have previously shown that passaging MCMV in human retinal pigment epithelial cells allows the virus to replicate to high titers in these cells due to the accumulation of adaptive mutations, such as loss-of-function mutations in the viral M117 gene. The M117 protein interacts with E2F transcription factors and activates E2F-dependent transcription. Here we show that activation of E2F3 is primarily responsible for MCMV’s inability to replicate in human cells. By transcriptome analysis, we identified two E2F3-induced serine proteases, FAM111A and FAM111B, as potential host restriction factors. By using shRNA-mediated gene knockdown and CRISPR/Cas9-mediated gene knockout, we demonstrated that FAM111B, but not its paralog FAM111A, suppresses MCMV replication in human and rhesus macaque cells. By immunofluorescence, we detected FAM111B predominantly in the nucleus of infected cells with enrichment in viral replication compartments, suggesting that it might play a role during viral replication. The fact that the FAM111B gene is conserved in primates but absent in rodents suggests that MCMV has not evolved to evade or counteract this restriction factor, which is not present in its natural host.

**Importance:** Viruses must counteract host cell defenses to facilitate viral replication. Viruses with a narrow host range, such as the cytomegaloviruses, are unable to counteract cellular defenses in cells of a foreign species. However, little is known about the cellular host range factors restricting cytomegalovirus replication. Here we show that MCMV induces the expression of the FAM111 proteases and that FAM111B, but not FAM111A that has previously been shown to restrict the replication of polyomavirus and orthopoxvirus host range mutants, acts as a cellular factor suppressing MCMV replication in human and rhesus monkey cells. The identification of FAM111B as a host range factor should provide new insight into the physiological functions of this poorly characterized protein.

## Introduction

Herpesviruses have coevolved with their respective hosts for millions of years and adapted to the host environment. One result of this adaptation is a restricted host range, a prominent feature of the cytomegaloviruses (CMV), representatives of the β-herpesvirus subfamily. They are highly species-specific as they replicate only in cells of their own or closely related host species (1). Human CMV (HCMV), an opportunistic pathogen causing serious disease in immunocompromised individuals, replicates in cells from humans or chimpanzees, but not in cells of more distant species such as mice or other rodents. Therefore, there is no small animal model to study HCMV pathogenesis. Experimental studies of CMV pathogenesis can only be conducted in animals by using their own CMV, for example by infecting mice with mouse CMV (MCMV) (2).

Early studies have shown that CMVs can enter non-permissive host cells and express a subset of viral genes, mainly of the immediate early (IE) class, but viral DNA replication and late gene expression are severely impaired. This has led to the conclusion that a post-penetration block to viral gene expression and replication is responsible for the species specificity of the CMVs (1, 3). Several studies suggested that the inability of MCMV to suppress cellular defenses in human cells plays an important role in the species specificity of this virus. One important restriction to MCMV replication in human cells is the inability of MCMV to inhibit apoptosis induced by viral DNA replication in human cells (4, 5). Inhibiting apoptosis in MCMV-infected human cells allows MCMV to replicate. However, viral replication under such conditions is not very efficient (4, 5), suggesting that additional checkpoints and restrictions exist. Another study provided evidence that MCMV can replicate in human cells with the help of some HCMV proteins, such as HCMV IE1 or pp71, that are capable of inhibiting human cellular defenses (6).

Subsequent studies on the mechanisms underlying the species restriction took advantage of MCMV mutants having spontaneously acquire the ability to replicate to high titers in human RPE-1 cells after a few passages (7). One set of adaptive mutations affected the MCMV M112-113 gene, which encodes the viral Early-1 (E1) proteins. The mutations increased the virus’ ability to disrupt PML bodies and form viral replication compartments (vRCs) (4). Of even greater impact were adaptive mutations in the MCMV M117 gene. We demonstrated that M117 interacts with E2F transcription factors and induces E2F-dependent transcription. This function is detrimental for MCMV replication in human cells (8), suggesting that human E2F target gene products might function as host restriction factors. Among the canonical members of the E2F transcription factor family, E2F1, E2F2, and E2F3 function as activators of gene transcription, whereas E2F4 and E2F5 are thought to act predominantly as transcriptional repressors. However, specific as well as redundant functions have been reported for each E2F (9, 10). For instance, while E2F1 and E2F3 are both necessary for cell cycle progression, E2F1 has been linked to the induction of apoptosis (11) whereas E2F3 regulates genes involved in DNA replication (9, 12)

In the present study, we found that E2F3-dependent transcription is particularly detrimental to MCMV replication in human cells. We identified an E2F-induced gene product, FAM111B, as a potent host restriction factor. Downregulation of FAM111B facilitated MCMV replication in human and rhesus cells. FAM111B, a serine protease of largely unknown function, accumulates in vRCs, suggesting a role of the protein during viral DNA replication.

## Results

### Activation of E2F3- but not E2F1-dependent transcription is detrimental for MCMV replication in human RPE-1 cells

In previous work, we demonstrated an interaction of the MCMV protein M117 with members of the E2F family of transcription factors in both murine and human cells (8). By using recombinant MCMVs expressing mutant M117 protein, we also demonstrated that the interaction with specific E2Fs can be modified. An N-terminal truncation mutant (M117-ΔNter) was unable to interact with E2Fs. Another mutant, M117-M4, retained the ability to interact with E2F3 and, to a lesser extent, with E2F1 (8). However, these specific interactions have been analyzed only in murine cells, and have not been verified in human cells. To confirm the specificity of these interactions in human cells, we infected human RPE-1 cells with recombinant MCMVs expressing Flag-tagged full-length (FL) M117, M117-ΔNter, or M117-M4 (Fig 1A). The interactions of WT and mutant M117 proteins with human E2Fs was analyzed by immunoprecipitation and immunoblotting. As shown in Fig 1B, M117-FL interacted with all endogenous human E2Fs tested, whereas no interaction with E2Fs was detected with the M117-ΔNter mutant. When M117-M4 was immunoprecipitated, only E2F3, but none of the other human E2Fs, was detected (Fig 1B).

**Figure 1.**
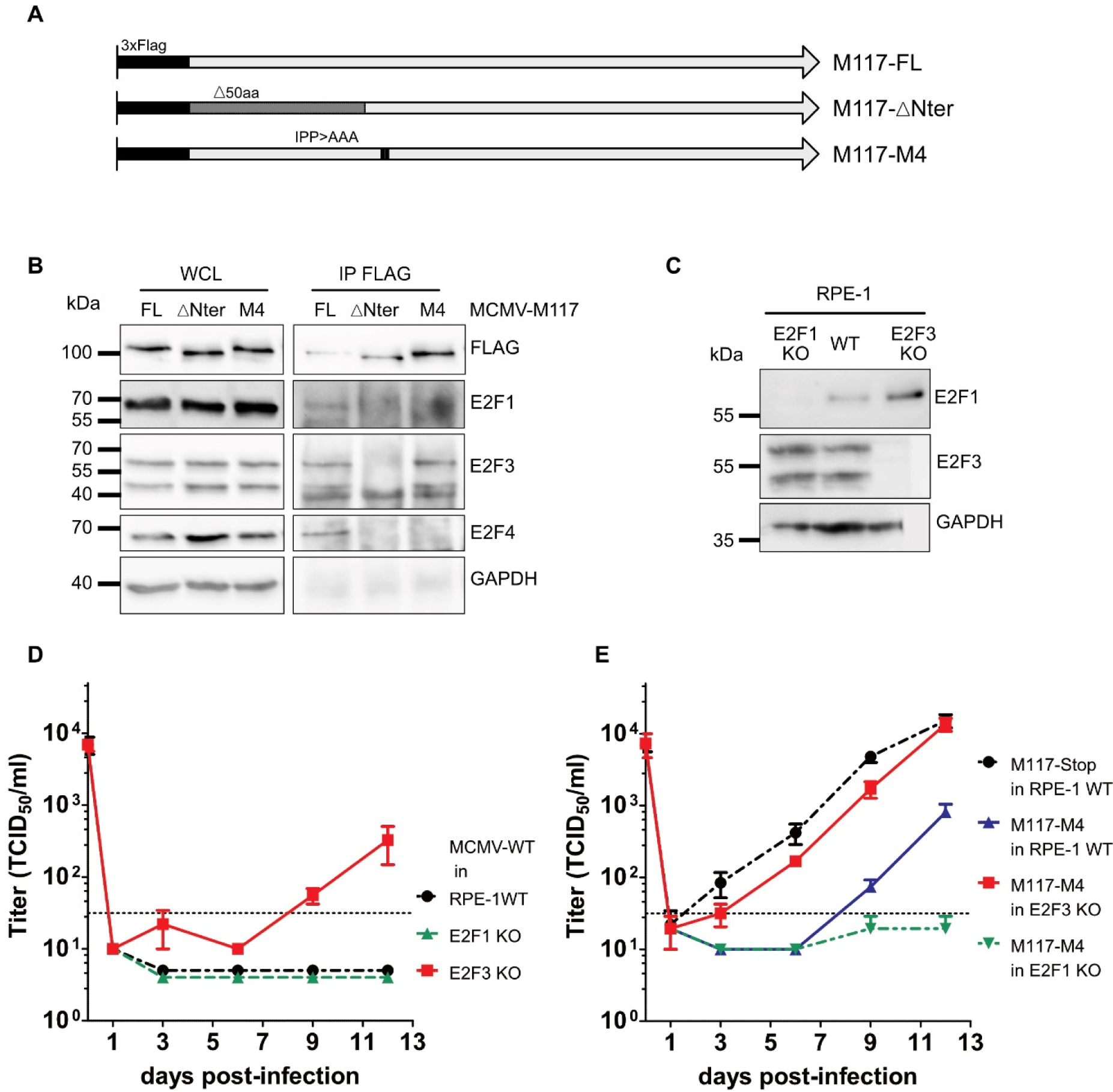
E2F3, but not E2F1, is detrimental for MCMV replication in human RPE-1 cells. (A) Schematic of the M117 mutants used in this study. FL, Full length; ΔNter, deletion of aa 1 to 50; M4, IPP>AAA substitution at positions 59–61. (B) RPE-1 cells were infected with MCMVs expressing 3xFlag-tagged M117-FL or mutant M117 proteins (MOI 2 TCID_50_/cell). Whole cell lysates (WCL) were collected 24 hpi and subjected to immunoprecipitation (IP) using an anti-Flag antibody. Co-precipitating proteins were detected by immunoblot analysis. (C) WCL of E2F1 and E2F3 KO RPE-1 cells were harvested and analysed by immunoblot. (D) Multistep replication kinetics of WT MCMV in WT RPE-1 and E2F KO cells. (E) Multistep replication kinetics of mutant MCMV strain in WT RPE-1 and E2F KO cells. Cells were infected at an MOI of 0.2 TCID_50_/cell. Virus released into the supernatant was titrated on murine fibroblasts. Mean ±SEM of triplicates are shown. DL, detection limit.

We previously showed that MCMV-M117-M4 replicated to lower titers than MCMV-M117-ΔNter in RPE-1 cells (8). As M117-M4 interacts only with E2F3, we hypothesize that E2F3 is a major restriction factor for MCMV in human cells. To evaluate the influence of E2F3 during MCMV replication, we generated E2F3 knockout (KO) RPE-1 cells by CRISPR-Cas9 gene editing. As a control, we also generated E2F1 KO RPE-1, as E2F1 and E2F3 have been shown to have partly redundant functions (9). The absence of the specific E2F proteins in the KO cells was verified by immunoblot analysis (Fig 1C). Next, both cell types were infected with MCMV to evaluate the importance of E2F3 and E2F1 in restricting MCMV replication. As shown in Fig 1D, MCMV replication in RPE-1 cells was detected only in E2F3 KO but not E2F1 KO cells. In the absence of E2F3, MCMV-M117-M4 replicated to similar titer as a virus lacking M117 (Fig 1E), indicating that the activation of E2F3 by M117 impairs MCMV replication in human cells. In contrast, the replication of MCMV-M117-M4 was impaired in the absence of E2F1 (Fig 1E), suggesting that E2F1 might have a more proviral function.

### M117 alters the transcriptome of MCMV-infected RPE-1 cells

E2Fs are transcription factors that regulate a large variety of genes involved in many different pathways (13). Due to the detected interaction of M117 with E2Fs and the observed replication of MCMV-ΔM117 in human cells, we hypothesized that M117 induces the expression of E2F-dependent genes restricting MCMV replication. Conversely, genes not induced by a mutant virus such as MCMV-ΔM117, are potential E2F-dependent restriction factors. In order to identify such E2F-dependent restriction factors, RPE-1 cells were synchronized by serum starvation for 72 hours before infection to enrich the proportion of cells in the same cell cycle stage (Fig 2A). Total RNA was extracted 24 hours post-infection (hpi) and subjected to RNA sequencing (RNA-seq) to compare the transcriptomes of WT MCMV and MCMV-ΔM117-infected cells (Fig 2B). The genes downregulated at least 2-fold in MCMV-ΔM117-infected RPE-1 cells compared to WT-infected RPE-1 cells are listed in Fig 2C. An analysis of 25 genes with a log_2_ score > 1.5 with Cytoscape (https://cytoscape.org/) showed that the E2F pathway was the main pathway differentially regulated by MCMV-ΔM117, thus confirming the regulation of E2Fs by M117.

**Figure 2.**
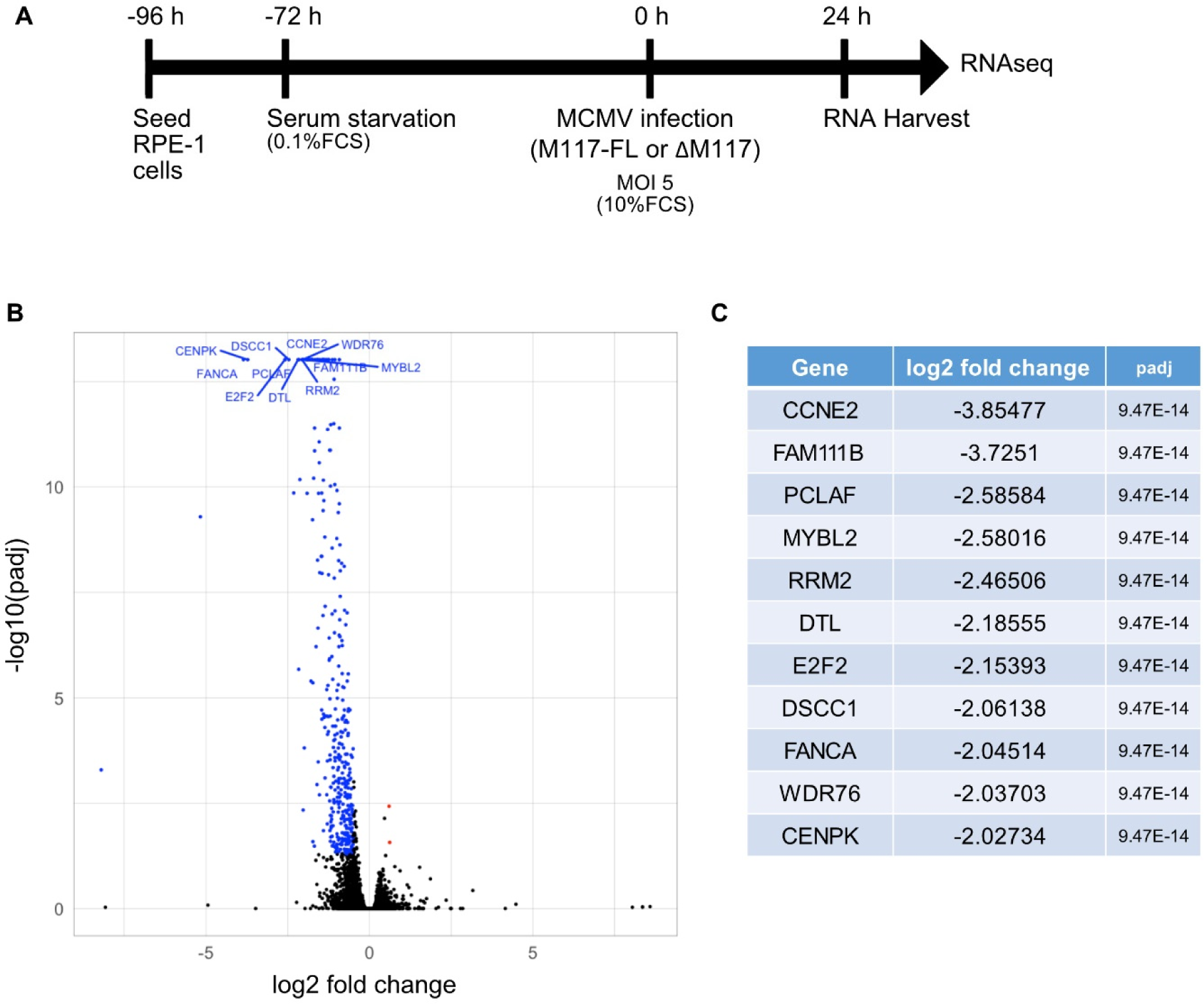
M117-dependent changes in the transcriptome of MCMV-infected RPE-1 cells. (A) Experimental setup for transcriptome analysis. (B) Volcano plot of the genes differentially expressed in infected RPE-1 cells (MCMV-ΔM117: MCMV-M117-FL). Blue dots represent genes expressed less in MCMV-ΔM117-infected cells. (C) List of genes downregulated at least 2-fold in MCMV-ΔM117-infected cells, with the adjusted p-value (padj). The data shown are the result of three biological replicates.

### CCNE2 is not an MCMV restriction factor

Cyclin E2 (CCNE2), a known E2F1 target (14, 15), was found to be the gene most strongly affected by the absence of M117 in MCMV-infected RPE-1 cells (Fig 2C). CCNE proteins are involved in cell cycle regulation as well as in DNA replication (16), but an antiviral function has not been described for these proteins. To test if CCNE2 is an MCMV restriction factor, we constructed recombinant MCMVs expressing CCNE2-specific shRNAs. To this end, the phosphoglycerate kinase (PGK) promoter within the previously described pReplacer plasmid (5) was substituted by a U6 promoter suitable for shRNA expression. Sequences encoding two *CCNE2*-specific shRNA (sh1 and sh2) and a scrambled (scr) control shRNA were inserted downstream of the U6 promoter. The plasmids were used to insert the U6 promoter and the shRNA sequences by homologous recombination into the nonessential m02-m06 locus of the MCMV-GFP BAC (Fig 3A). The recombinant BACs were sequence-verified and used to reconstitute infectious viruses. In RPE-1 cells infected with the control virus (MCMV-scr), we detected an upregulation of *CCNE2* mRNA levels by quantitative RT-PCR. In contrast, both viruses encoding *CCNE2*-specific shRNAs induced lower *CCNE2* mRNA levels compared to MCMV-scr (Fig 3B), confirming that the shRNAs are functional. Next, we assessed the ability of these viruses to replicate in human RPE-1 cells. None of the viruses replicated in RPE-1 cells as determined by multistep replication kinetics (Fig 3C), suggesting that the downregulation of CCNE2 is not sufficient to allow MCMV replication in human RPE-1 cells. However, we did observe the appearance of small foci of infected cells with MCMV-shRNA2 (Fig 3D) but not with the control viruses nor with MCMV-shRNA1. Hence, it remains unclear whether this minor phenotype is a *CCNE2*-specific or an off-target effect of *CCNE2* shRNA2.

**Figure 3.**
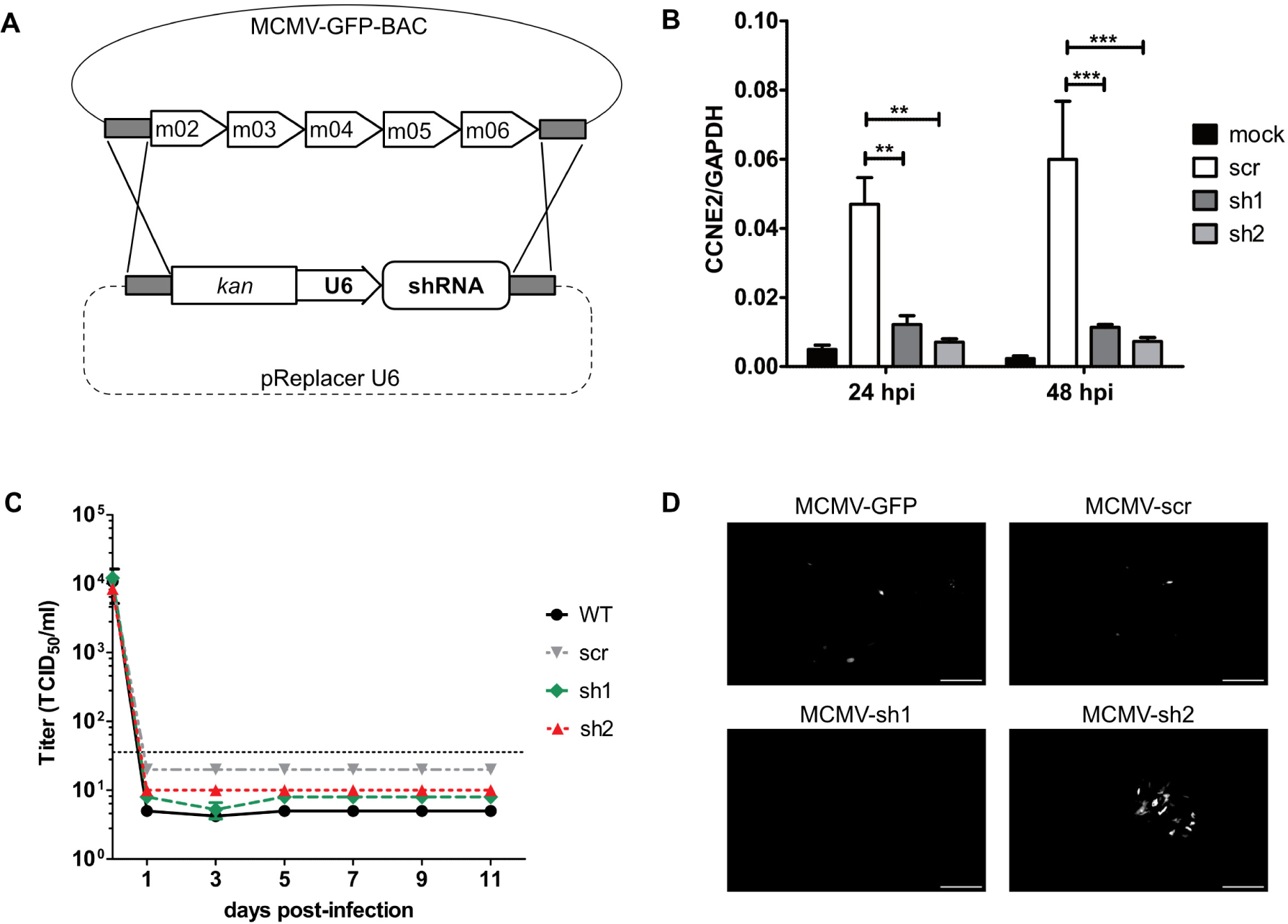
CCNE2 is not an MCMV restriction factor. (A) Construction of recombinant MCMV BACs. Homology arms are shaded grey, shRNAs are driven by a U6 promoter. Kan, kanamycin resistance marker. (B) RPE-1 cells were infected with MCMVs expressing CCNE2-specific shRNAs (sh1 and sh2) or scrambled (scr) control shRNA (MOI 3 TCID_50_/cell). RNA was extracted at 24 and 48 hpi and analyzed by quantitative RT-PCR. Mean ± SEM of three independent experiments are shown. (C) Multistep replication kinetics of WT (MCMV-GFP) and recombinant MCMVs in RPE-1 cells (MOI 0.2). Virus released into the supernatant was titrated on murine fibroblasts. Mean ±SEM of triplicates are shown. DL, detection limit. (D) Fluorescence images of cells infected for 9 days (as in C). Scale bar, 500 µm.

### E2F3 is a major regulator of FAM111A and FAM111B during MCMV infection

As CCNE2 was ruled out as a major restriction factor, we next focused on FAM111B, the second gene in the list (Fig 2C). FAM111B is a trypsin-like cysteine/serine peptidase conserved in primates but absent in rodents (17). Interestingly, FAM111B has been described as a restriction factor for human adenovirus type 5 (Ad5) (18) whereas its paralog, FAM111A is a host range factor for SV40 and orthopoxviruses (19, 20). We therefore included FAM111A in our analysis, as it is also downregulated in our transcriptome analysis, but with a lower log_2_ fold change of −0.92.

All previous studies involving FAM111A and FAM111B were performed in cancer cells, which have a dysregulated cell cycle. In our study, we used RPE-1 cells, which were immortalized with human telomerase reverse transcriptase (hTERT) but are not transformed. Therefore, we tested whether FAM111A and FAM111B are expressed in a cell cycle-dependent manner in RPE-1 cells, similarly to what was described for other cell types (21, 22). The peak levels of FAM111A and FAM111B in RPE-1 cells were detected during the S phase (Fig 4 A and B). Moreover, FAM111A and FAM111B levels were reduced upon treatment with HLM006474, a Pan-E2F inhibitor (23) (Fig 4C). These results confirmed the E2F-dependent expression of these proteins.

**Figure 4.**
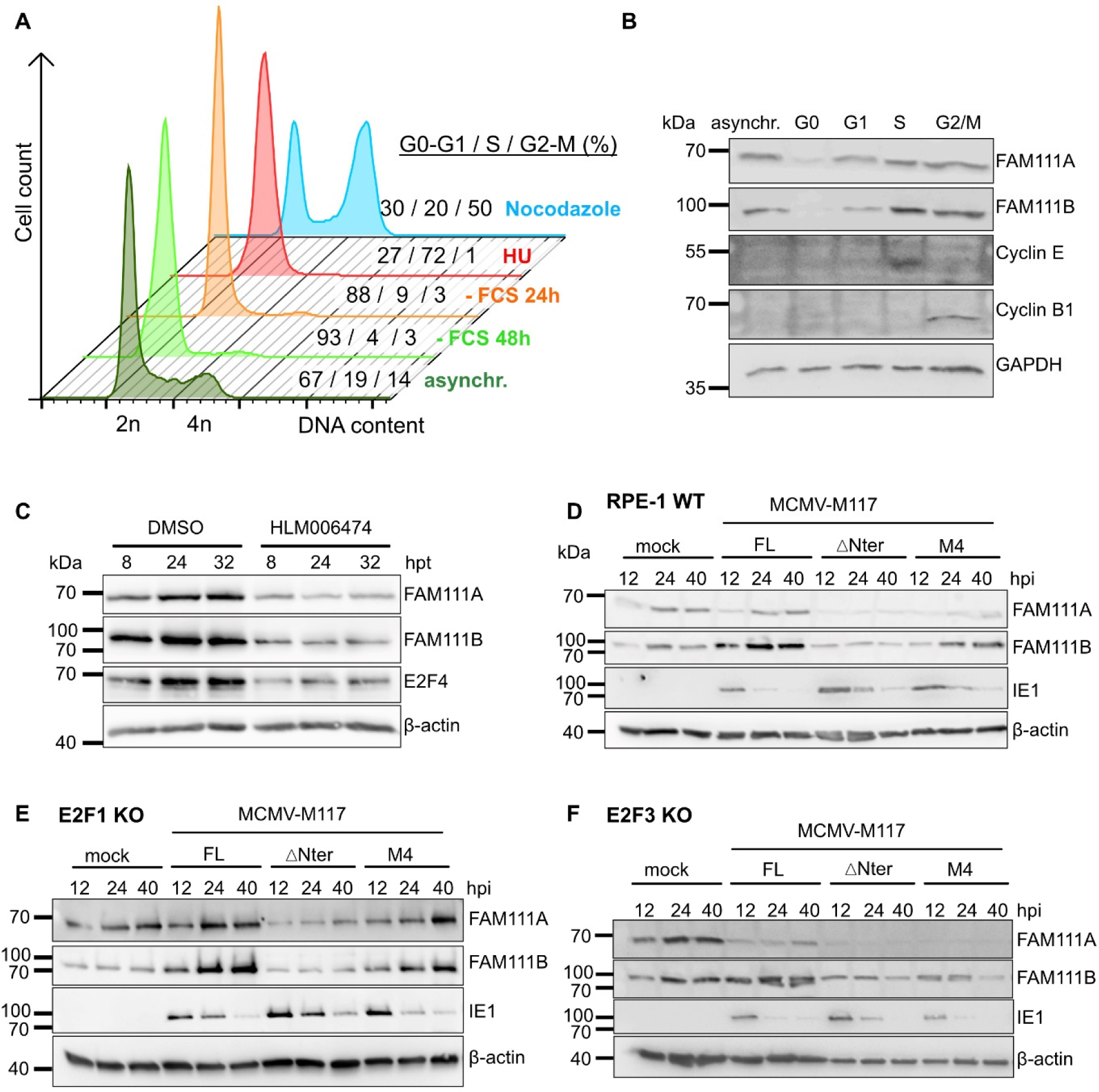
E2F3 is a major regulator of FAM111A and FAM111B during MCMV infection. (A) RPE-1 cells were arrested in G0 by contact inhibition and 48 h serum starvation (-FCS 48 h), in G1 by 24 h serum starvation (-FCS 24 h), in S phase by 24 h treatment with 1 mM hydroxyurea (HU) and in G2/M by 24 h treatment with 200 ng/ml nocodazole. Asynchronous (asynchr.) cells are shown for comparison. The cells were fixed and stained with propidium iodide. The DNA content was analyzed by flow cytometry to verify synchronization. Cells in G0 and G1 have a DNA content of 2n, cells in S phase between 2n and 4n, and cells in G2/M a 4n DNA content. The percentages of cells in each cell cycle phase are indicated. (B) FAM111A and FAM111B expression in cell-cycle arrested cells was analyzed by immunoblot. Cyclin E and cyclin B1 were analyzed as markers for S and G2/M phases, respectively. GAPDH was used as a loading control. (C) Subconfluent RPE-1 cells were treated with an E2F inhibitor (HLM0006474, 40 μM) or vehicle (DMSO). Cell lysates were harvested at the indicated times (hours post-treatment, hpt) and analyzed by immunoblot. (D to F) WT RPE-1 cell, E2F1 KO, and E2F3 KO cells were synchronized by serum starvation for 24 h and infected with MCMVs expressing full-length or mutant M117 (MOI 5). Cell lysates were harvested at the indicated times post-infection and analyzed by immunoblot. The MCMV immediate-early 1 (IE1) protein and β-actin were used as infection and loading controls, respectively. Representative blots out of multiple experiments are shown. hpt: hours post treatment.

Next, we wanted to test whether MCMV regulates the expression of FAM111A and B in an M117-dependent manner. We synchronized RPE-1 cells in the G1 phase and infected them with different M117-mutated MCMVs. Under these conditions, MCMV-M117-FL strongly increased the expression of FAM111B whereas the expression of FAM111A was only slightly changed. In contrast, MCMV-M117-ΔNter did not induce the expression of the two proteins (Fig 4D), suggesting that M117 regulates FAM111B expression. MCMV-M117-M4 also induced the expression of FAM111A and FAM111B (Fig 4D), suggesting that the induction was mediated by E2F3. A previous study suggested that E2F1 is the regulator of FAM111B (18). However, in E2F1 KO RPE-1 cells the regulation of FAM111A and B by MCMV did not differ from WT cells (Fig 4E), suggesting that the regulation of FAM111A and FAM111B in MCMV-infected RPE-1 cells is not dependent on E2F1. In contrast, the upregulation of FAM111A and B by both MCMV-M117-FL virus and MCMV-M117-M4 was completely abrogated in E2F3 KO RPE-1 cells, resulting in similar expression levels as the ones observed in mock-infected cells or cells infected with MCMV-M117-ΔNter (Fig 4F). These data strongly suggested that E2F3 is the major regulator of FAM111A and FAM111B in MCMV-infected RPE-1 cells.

### FAM111B but not FAM111A is an MCMV host restriction factor in primate cells

FAM111A and FAM111B are paralogous proteins with overlapping functions (24, 25). However, in their function as viral restriction factors both proteins appear to be more specific. While FAM111A is a restriction factor for host range mutants of SV40 and orthopoxviruses (19, 20), FAM111B is a restriction factor for Ad5 (18). To identify whether one of these two proteins functions as an MCMV restriction factor, we generated recombinant MCMVs encoding shRNAs specific for FAM111A (MCMV A-sh) or FAM111B (MCMV B-sh), using the same method as described above for CCNE2 (Fig 3A). Two different shRNA sequences were used for FAM111A (Ash1 and Ash2) and FAM111B (Bsh1 and Bsh2). Infection of RPE-1 cells with MCMV-Ash1 or MCMV-Ash2 resulted in a downregulation of FAM111A but not FAM111B, and infection with MCMV-Bsh1 or MCMV-Bsh2 resulted in a downregulation of FAM111B but not FAM111A (Fig 5A), indicating a high specificity of the shRNA sequences used. In multistep replication kinetics, only the knockdown of FAM111B allowed MCMV replication in RPE-1 cells, whereas the knockdown of FAM111A had no impact (Fig 5B). Similar results were obtained with two other human epithelial cell types, ARPE-19 retinal epithelial cells and RPTEC kidney epithelial cells (Fig 5C and 5D), indicating that the phenotype is not specific to RPE-1 cells used in the previous experiments.

**Figure 5.**
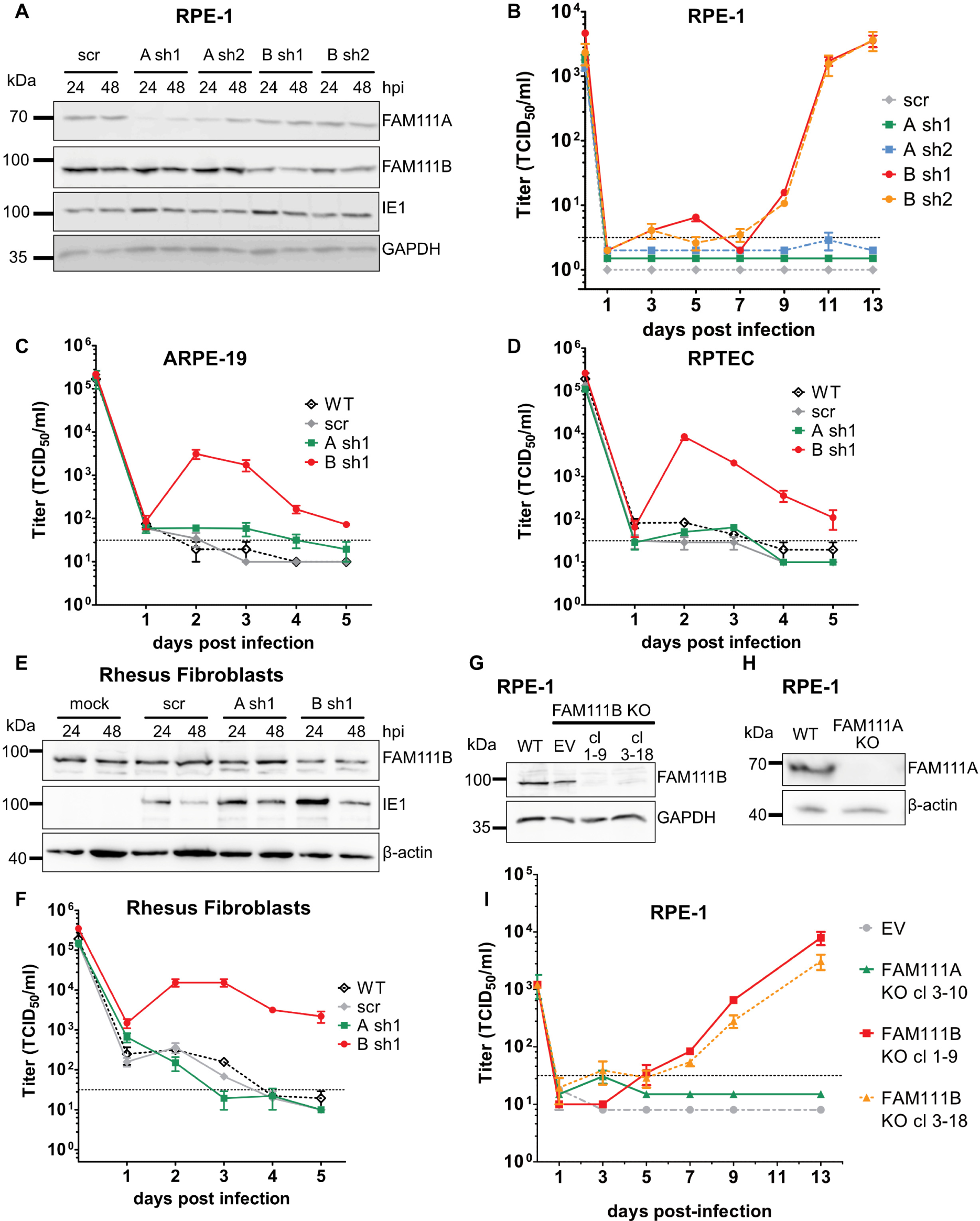
FAM111B, but not FAM111A, is an MCMV host restriction factor in primate cells. (A) RPE-1 cells were infected (MOI 5 TCID_50_/cell) with recombinant MCMV-GFP encoding FAM111A-specific (A sh1, A sh2), FAM111B-specific (B sh1, B sh2), or scrambled (scr) control shRNA. Cell lysates were harvested 24 and 28 hpi and analyzed by immunoblot. (B) RPE-1 cells (MOI 0.2), (C) ARPE-19 (MOI 1) and (D) RPTEC (MOI 1) were infected with the same viruses as above. Viral replication kinetics were determined by titration of virus released into the supernatant. (C) Rhesus fibroblasts were infected (MOI 1.5 TCID_50_/cell) with the same viruses as above and analyzed by immunoblot or infected at MOI 1 TCID_50_/cell (D) with the same viruses as above. Viral replication kinetics were determined by titration of virus released into the supernatant. Mean ±SEM of triplicates are shown. DL, detection limit. (E, F) Cell lysates of WT RPE-1 cells, empty vector (EV)-transduced RPE-1 cells, two FAM111B KO RPE-1 clones (cl 1-9 and 3-18), and one FAM111A KO RPE-1 (cl 3-10) clone were analyzed by immunoblot. (G) Multistep replication kinetic of WT MCMV in the same RPE-1 as in E. Supernatants of infected cells were harvested at the indicated times post infection and titrated. Mean ±SEM of triplicates are shown. DL, detection limit.

Due to the conservation of FAM111A and FAM111B in primate cells, we could use the Ash1 and Bsh1 sequences to downregulate the homologous genes in cells of rhesus macaques. We were unable to detect FAM111A in rhesus fibroblasts by immunoblot with an antibody against human FAM111A, but we were able to detect FAM111B and could confirm the downregulation of the rhesus FAM111B protein level by MCMV-Bsh1 (Fig 5E). When we analyzed viral replication kinetics in rhesus fibroblasts, we found that neither WT MCMV nor MCMV-scr replicated in these cells (Fig 5F). MCMV-Ash1 did not replicated in rhesus fibroblasts either, but MCMV-Bsh1 did replicate (Fig 5F), indicating that a knockdown of FAM111B facilitates MCMV replication in these cells.

To confirm the antiviral function of FAM111B in RPE-1 cells, we generated human RPE-1 cells in which FAM111B was knocked out by CRISPR-Cas9 gene editing. As a control, we used FAM111 KO cells generated by the same method. FAM111B protein was not detected in two FAM111B KO RPE-1 clones (1-9 and 3-18) obtained by using two independent guide RNAs (Fig 5G). Similarly, FAM111A expression was not detected in FAM111A KO RPE-1 cells (Fig 5H). When we analyzed MCMV replication kinetics in these cells, we detected viral replication to high titer only in the absence of FAM111B but not in the absence of FAM111A (Fig 5I). Altogether, these data indicated that FAM111B, but not FAM111A, is a conserved host restriction factor for MCMV in primate cells.

### FAM111B antiviral function requires human cellular context

The FAM111B gene is conserved in the genomes of primates but it is not present in rodent genomes (17). Therefore, we wondered whether human FAM111B can inhibit MCMV replication when expressed in mouse cells as it does in human cells. To test this, we generated recombinant MCMVs expressing either HA-tagged WT FAM111B or one of two mutant forms. The S628N mutation, present in the hereditary fibrosing poikiloderma syndrome, leads to a hyperactive form of the protein, whereas the D544N mutation inactivates the FAM111B protease activity (25). WT and mutant FAM111B cDNAs, driven by a phosphoglycerate kinase (PGK) promoter, were inserted into the non-essential m02-m06 locus of MCMV by BAC recombineering. Transfection of the recombinant MCMV BACs into murine fibroblasts resulted in reconstitution of infectious virus and plaque formation, suggesting that the inserted FAM111B gene did not preclude MCMV replication.

To verify FAM11B expression, we analyzed infected cells by immunoblot analysis. As mouse cells do not express endogenous FAM111B, the protein was not detected in mock-infected and WT MCMV-infected cells (Fig 6A). However, upon infection with the recombinant viruses, FAM111B was detected with antibodies directed against the HA-tag or the protein itself (Fig 6A). The protein levels of FAM111B S628N appeared to be somewhat lower, possibly due to an increased autocleavage of the hyperactive protein (25).

**Figure 6.**
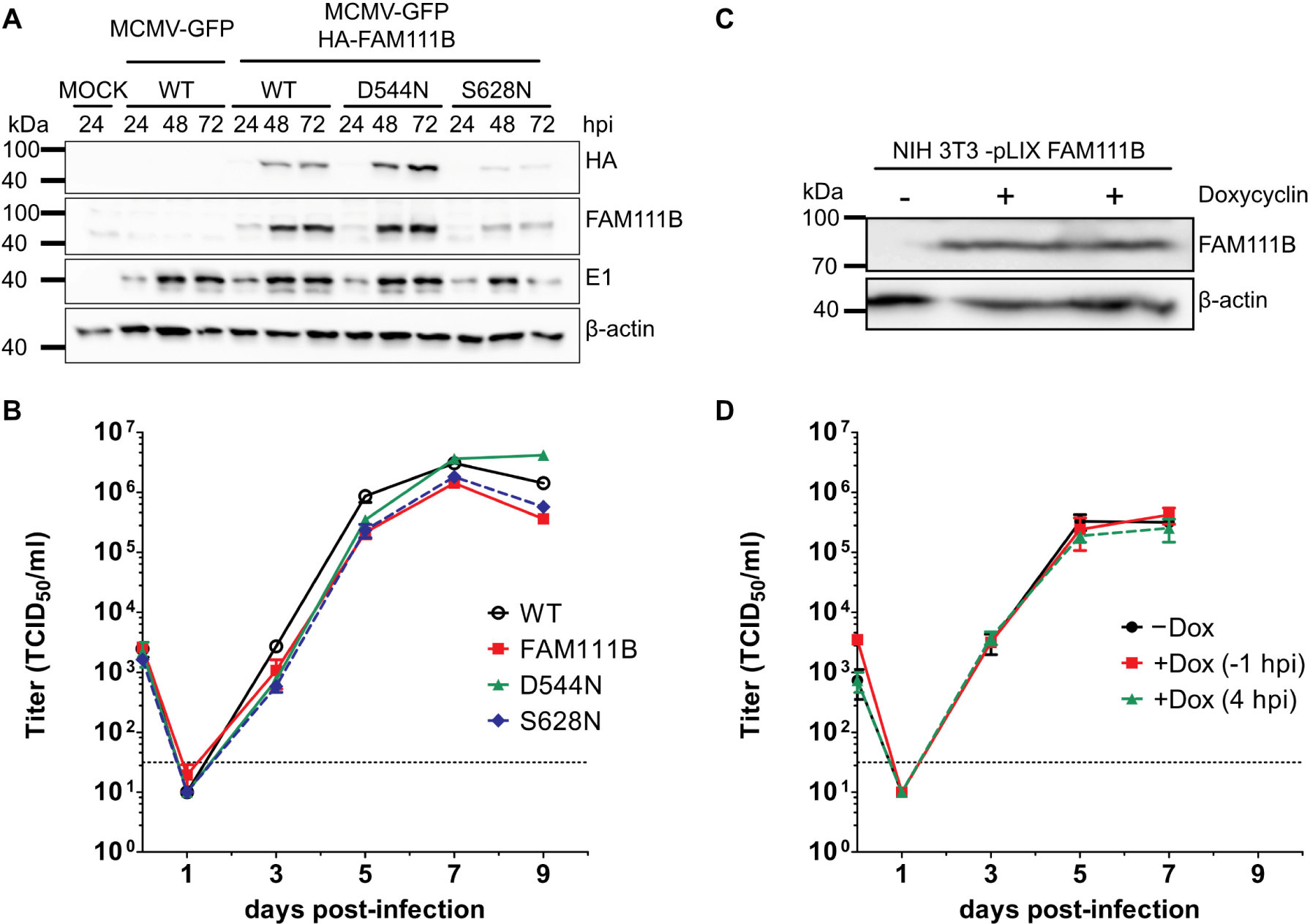
FAM111B antiviral function require human cellular context. (A) 10.1 mouse fibroblasts were infected (MOI 1 TCID_50_/cell) with recombinant MCMVs encoding HA-tagged WT or mutant FAM111B. Cell lysates were analyzed by immunoblot. (B) 10.1 cells were infected (MOI 0.02) with the same viruses as above to analyze multistep replication kinetics. Mean ±SEM of triplicates are shown. DL, detection limit. (C) NIH-3T3 fibroblasts transduced with a lentiviral vector encoding tet-inducible FAM111B were induced with 2 µg/ml doxycyline or left untreated. Cell lysates were harvested and analyzed by immunoblot. (D) Transduced NIH-3T3 cells were treated with doxycycline 1h before (−1 hpi) or 4 h after (+4 hpi) infection with WT MCMV (MOI 0.02) to analyze multistep replication kinetics. Mean ±SEM of triplicates are shown.

To test whether FAM111B expression inhibits MCMV replication in murine cells, we analyzed the replication kinetics of the viruses described above. No significant differences in the replication of the FAM111B-expressing recombinant viruses compared to WT MCMV were observed (Fig 6B), suggesting that FAM111B does not restrict MCMV replication in murine cells. To rule out the possibility that the expression of FAM111B by MCMV is temporally inadequate for its antiviral function, we also generated transduced murine fibroblasts with a lentiviral vector expressing FAM111B in a doxycycline-inducible fashion (Fig 6C). In these cells, MCMV replication was unaffected, no matter whether FAM111B expression was induced before (−1 hpi) or after MCMV infection (+4 hpi) (Fig 6D). These results confirmed that FAM111B does not function as an MCMV restriction factor in murine cells.

To analyze the intracellular distribution of FAM11B, we generated a recombinant virus expressing an eGFP-tagged FAM111B, which allowed us to visualize the localization of FAM111B in MCMV-infected murine fibroblasts. While FAM111B could not be detected in mock-infected murine cells (Fig 7A), the protein was detected throughout the cytoplasm and nucleus of infected cells (Fig 7A). FAM111B accumulation in punctate structures within the nucleus was observed, but these structures did not colocalize with the viral E1 protein, a marker of vRCs. Unfortunately, the infection of human RPE-1 cells with MCMV-eGFP-FAM111B resulted in a low eGFP signal, suggesting that overexpression of FAM111B is not well-tolerated in these cells. However, we could detect endogenous FAM111B by immunofluorescence staining with a FAM111B-specific antibody. FAM111B was not detected in FAM111B KO RPE-1 cells, confirming the specificity of the immunostaining. In both, non-infected and MCMV-infected WT RPE-1 cells, FAM111B had a dispersed nuclear localization (Fig 7B). In MCMV-infected cells, FAM111B accumulated in large vRCs as indicated by colocalization with the viral E1 protein at late times post-infection. Taken together, these data suggest that function and localization of FAM111B are altered in murine cells. Additional human-specific factors are probably required for its proper localization and accumulation in vRCs where it might exert its antiviral function.

**Figure 7.**
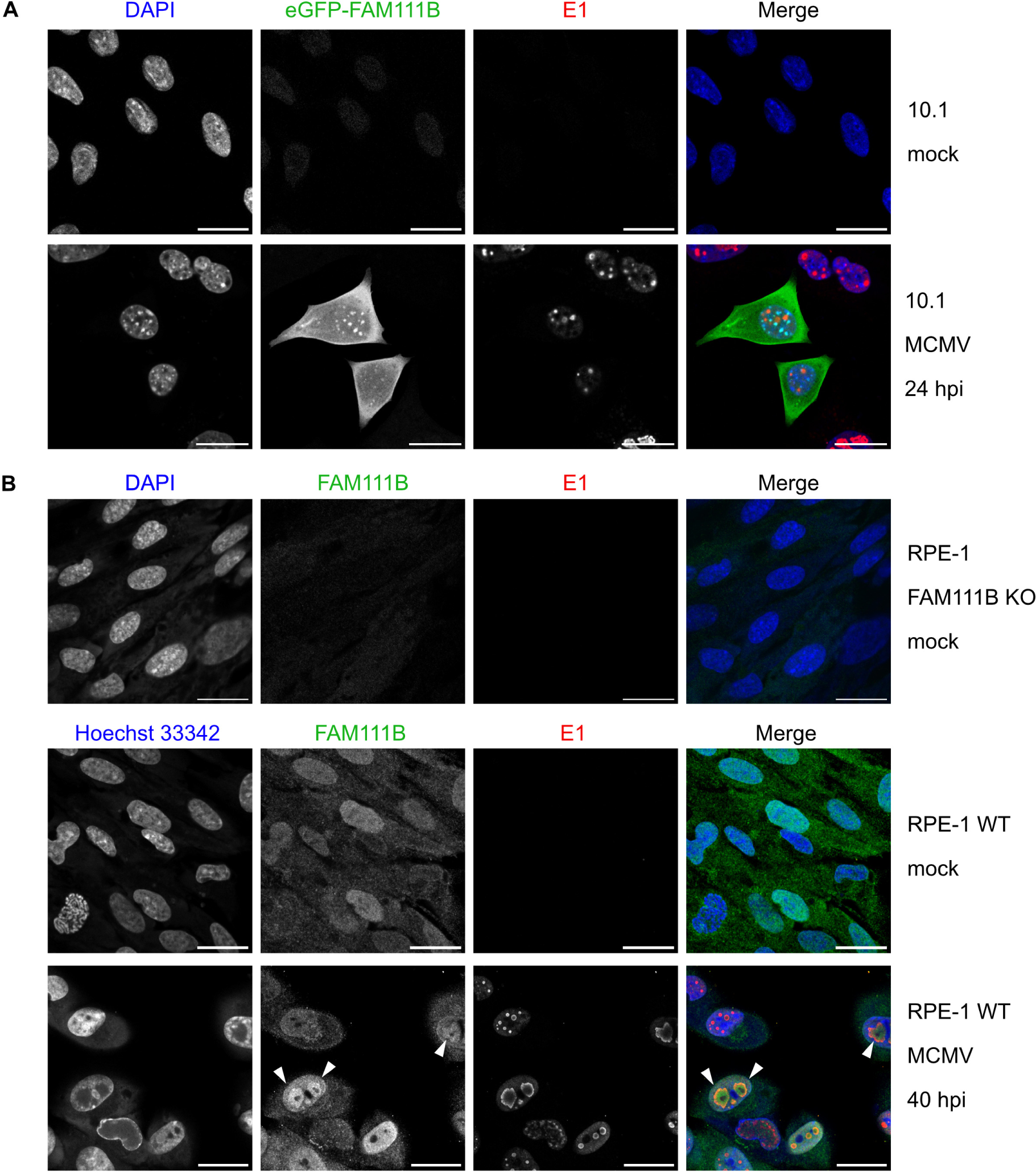
FAM111B localizes to the viral replication compartments in human cells. (A) Immunofluorescence of 10.1 fibroblasts mock-infected or infected with MCMV-eGFP-FAM111B (MOI 1 TCID_50_/cell). Cells were fixed 24 hpi and stained with an antibody recognizing the MCMV E1 protein (red), a marker for vRCs. Nuclei were stained with DAPI. Scale bar, 25 µm. (B) Immunofluorescence of FAM111B KO or WT RPE-1 cells mock-infected or infected with WT MCMV (MOI 1 TCID_50_/cell). Cells were fixed 40 hpi and stained with antibodies recognizing FAM111B (green) or MCMV E1 (red). Nuclei were stained with DAPI or Hoechst 33342. Arrow heads indicate FAM111B in vRCs. Scale bar, 25 µm.

## Discussion

In this study, we show that the MCMV M117 protein interacts with human E2F transcription factors and that activation of the E2F3, but not E2F1, is detrimental to MCMV replication in human cells: MCMV was able to replicate in E2F3 KO, but not in E2F1 KO cells (Fig 1D). These results suggest that E2F3 is required for the upregulation of host factors inhibiting MCMV replication. However, since the E2F transcription factors have partly redundant functions (26), a role of other E2Fs in restricting MCMV cannot be ruled out.

We hypothesized that cellular genes upregulated in an M117-dependent fashion in MCMV-infected human cells might encode inhibitory host factors. We tested the top two hits of a transcriptomic analysis and identified FAM111B as a major restriction factor for MCMV. Knockdown of FAM111B expression by shRNA or knockout by CRISPR/Cas9 gene editing was sufficient to facilitate MCMV replication in human epithelial cells and rhesus fibroblasts. In contrast, knockdown or knockout of the closely related FAM111A protein did not facilitate MCMV replication in these cells, even though FAM111A was previously shown to function as a restriction factor of SV40 and poxvirus host range mutants.

FAM111A is a trypsin-like serine protease whose function remains incompletely understood (17, 27). Several studies demonstrated its involvement in DNA replication. FAM111A was shown to localize at nascent DNA and promote DNA replication by interacting with proliferating cell nuclear antigen (PCNA) (28). It also mitigates the effect of protein obstacles on replication forks through its protease domain, thereby inhibiting replication fork stalling (29). The *FAM111B* gene is a paralog of *FAM111A* located adjacent to *FAM111A* on chromosome 11, suggesting that FAM111B arose by gene duplication. To date, little is known about the function of the FAM111B protein (17, 27, 30). Transcriptomic and proteomic interaction studies suggested a role of FAM111B in cell cycle regulation, DNA damage repair and replication, and apoptosis. However, further studies will be necessary to unravel the specific functions of FAM111B in these processes.

In 2012, Fine *et al*. demonstrated that FAM111A functions as a restriction factor for an SV40 host range mutant lacking the C-terminus of the large T (LT) antigen (19). In WT SV40-infected cells, the C-terminal part of LT interacts with FAM111A and blocks its antiviral activity. A later study showed that FAM111A relocalizes to SV40 vRCs in the host cell nucleus where it is thought to cleave factors necessary for viral replication (21). FAM111A is also a restriction factor of poxviruses, large DNA viruses replicating in the cytoplasm (20). It inhibits the replication of host range mutants lacking the viral serine protease inhibitor SP-I. FAM111A inhibits poxvirus replication by targeting the viral DNA-binding protein I3, an essential component of vRCs, for degradation by autophagy (31). Moreover, FAM111A degrades the nuclear pore complex through its protease activity (32). SP-1 counteracts this activity by interacting with FAM111A and preventing its nuclear export (31).

The first evidence that FAM111B might also function as a host restriction factor was provided in a recent study by Ip *et al*. (18). The study revealed that human Ad5 first upregulates FAM111B transcription by its E1A protein. At later times post-infection, the viral E1B-55k and E4orf6 proteins are responsible for decreasing FAM111B protein levels. Knockdown of FAM111B in human A549 cells slightly increased viral titers, suggesting that FAM111B might be detrimental to Ad5 replication (18). In contrast, MCMV replication is strongly affected by the presence or absence of FAM111B, as we show here. MCMV cannot replicate in human or rhesus cells but replicates when FAM111B expression is reduced or abolished. Hence, FAM111B is an MCMV restriction factor. Elucidating the mechanism by which FAM111B inhibits MCMV replication remains a difficult task as long as so little is known about the physiological functions of this protein. However, the accumulation of FAM111B in MCMV vRCs (Fig. 7) suggests an inhibitory function within vRCs, possibly by cleavage of vRC components as it has been proposed for FAM111A in SV40 vRCs (21).

The question of why MCMV cannot antagonize the antiviral activity of FAM111B has an obvious answer: the mouse does not possess a FAM111B gene (17). Therefore, there has never been a need for MCMV to antagonize this protein. A more interesting question is whether FAM111B can inhibit the replication of HCMV and other human herpesviruses. It is conceivable that human herpesviruses have evolved to circumvent or block the antiviral activity of FAM111B. Identifying viral FAM111B antagonists might also provide new insights into the physiological function of this enigmatic protein.

## Material and Methods

### Cell culture and viruses

Human telomerase reverse transcriptase (hTERT) immortalized human retinal pigment epithelial cells (hTERT RPE-1, ATCC CRL-4000), rhesus fibroblasts (33), mouse NIH-3T3 (ATCC CR1658) fibroblasts, and mouse 10.1 fibroblasts (34) were maintained in complete Dulbecco’s modified Eagle medium (DMEM) supplemented with 10% fetal calf serum (FCS) and 100 IU penicillin/100 µg streptomycin (P/S) at 37°C and 5% CO_2_. ARPE-19 retinal pigment epithelial cells (ATCC CRL-2302) were maintained in DMEM/Ham’s F12 (1:1) medium supplemented with 10% FCS and P/S. RPTEC/TERT1 (ATCC CRL-4031) were grown in DMEM/Ham’s F12 (1:1) supplemented with P/S, 2% FCS, 36 ng/mL hydrocortisone, 10 ng/mL EGF and 1x insulin/transferrin/selenium.

The GFP-expressing (MCMV-GFP) and the “repaired” WT MCMV Smith strain (pSM3fr-MCK-2fl) strain have been described previously (35, 36). Recombinant MCMVs expressing Flag-tagged WT and mutant M177 have been described (8). All viruses were propagated in 10.1 fibroblasts. For high-MOI infections, centrifugal enhancement of infection (1000 × *g*, 30 min) was used (2).

### Viral replication kinetics

Cells were seeded at low density on the day prior to infection in six-well dishes. The infection was performed at a low MOI (MOI<1). Input virus was removed after 4h and fresh medium was added. Supernatants were harvested at different times post-infection and stored at −80°C. Titration on 10.1 cells was done using the median tissue culture infective dose (TCID_50_) method (37). For the MCMV-shCCE2 replication kinetic, fluorescence imaging was performed using a Nikon Eclipse Ts2 microscope. Image analysis was done with ImageJ software (38).

### BAC mutagenesis

The introduction of FAM111B or the shRNAs into the nonessential m02-m06 region of MCMV-GFP was done by using the pReplacer plasmid (5) as described below. The integrity of the mutant BACs was verified by restriction fragment length pattern analysis and sequence analysis of the mutated region. To reconstitute infectious virus from MCMV BACs, purified BAC DNA was transfected into 10.1 fibroblasts using Polyfect (Qiagen).

### Plasmids

The U6 promoter and the shRNA-scr sequence were PCR-amplified from pLKO.1 shRNA scr (39) using a forward primer containing a MfeI restriction site (TATA<u>CAATTG</u>GAGGGCCTATTTCCCATGATT) and a reverse primer containing a BamHI restriction site (TTAA<u>GGATCC</u>TGCCATTTGTCTCGAGGTC). The PCR product was cleaved with MfeI and BamHI and inserted into the pReplacer plasmid (5) digested with EcoRI and BamHI in order to generate pReplacer-U6-shscr. The cloning of specific shRNA sequences into pReplacer-U6 was performed as described (www.addgene.org/protocols/plko/) using the following siRNA sequences: CCNE2 sh1 (GCTCTTAAAGATGCTCCTAAA), CCNE2 sh2 (CCAGACACATACAAACTATTT), FAM111A sh1 (GTCAATGTGTAAGGGTGACAT), FAM111A sh2 (GCATCGACTGAATGTGTCAAA), FAM111B sh1 (GCCTGCCTAGTGATTCTCATT) and FAM111B sh2 (GCGAACAGCTTACATATTATA). The FAM111B cDNA was amplified with primers TATACTGCAGATGTACCCATACGACGTCCCAGACTACGCTAATTCCATGAAG-ACTGAAGAAAAC and TTAACTGCAGCTAACATTCCATGGGTTCAATC) from cDNA obtained from RPE-1 cells and cloned into pReplacer or the tet-on lentiviral vector pLIX-402 (Addgene #41394) using PstI restriction sites. The HA-tag sequence was included in the forward primer. FAM111B mutations S628N and D544N were introduced into pReplacer-HA-FAM111B using NEBuilder HiFi DNA Assembly Master Mix (NEB). The pReplacer-eGFP-FAM111B was generated by inserting FAM111B into peGFP-C1 (Clontech) with EcoRI and BamHI. eGFP-FAM111B was subcloned in pReplacer using NheI and BamHI restriction enzymes.

### Generation of knockout RPE-1 cells

The lentiviral CRISPR/Cas9 vector pSicoR-CRISPR-PuroR was used to generate the different KO clones essentially as described (40, 41). The following guideRNAs were designed using the E-CRISPR design tool (www.e-crisp.org) and cloned individually into the lentiviral vector: E2F3, GCTGCAGTCTGTCTGAGGAT; E2F1, GCCACAGGTGAAGCGGAGGC; FAM111A, GGCCAAGAAATGCTTGTGCG; FAM111B g1, GCGCTATGGAAGATGACCAG and FAM111B g3, GGAGTTTCCATTCAACGTAA.

### Lentiviral transduction

Lentiviruses were generated using standard third-generation packaging vectors in HEK-293T cells as described (42). For generating KO cells, RPE-1 cells were transduced in the presence of polybrene (5 µg/mL) (Sigma) with lentiviral CRISPR/Cas9 vectors containing gRNA or empty vector (EV). The cells were selected with 2.5 µg/mL puromycin (Sigma) and subcultured to obtain single cell clones for each gRNA. Gene knock-out cell clones were verified by screening for the absence of the relevant protein by immunoblot analysis. To generate NIH-3T3 cells expressing FAM111B in a tet-inducible fashion, NIH-3T3 cells were transduced with pLIX-FAM111B and selected with 2 µg/mL puromycin. Expression of FAM111B was induced by adding 2 µg/mL doxycyxline (Biomol).

### Quantitative real-time PCR (qPCR)

To quantify cyclin E2 transcripts, RPE-1 cells were infected at an MOI of 3. Total RNA was extracted from infected cells at 24 and 48 hpi using an innuPREP RNA Mini kit (Analytik Jena). Contaminating DNA was removed using a TURBO DNA-free kit (Ambion). cDNA was synthesized from 1 µg of the extracted RNA by using the RevertAid H Minus Reverse Transcriptase, oligo-dT primers, and the RNase inhibitor RiboLock (Thermo Fisher Scientific). qPCR was done with a QuantStudio3 Real Time PCR System (Thermo Fisher Scientific) using 10 ng of cDNA, the PowerTrack SYBR Green Mastermix (Thermo Fisher Scientific). The following primers were used: *GAPDH* (CCCACTCCTCCACCTTTGACG and GTCCACCACCCTGTTGCTGTAG); *CCNE2* (CTTACGTCACTGATGGTGCTTGC and CTTGGAGAAAGAGATTTAGCCAGG). *CCNE2* transcripts levels were normalized to the housekeeping gene glyceraldehyde-3-phosphate dehydrogenase (*GAPDH)*.

For MCMV genome quantification, total DNA was extracted from MCMV-infected RPE-1 cells using an InnuPREP DNA Mini Kit (Analytik Jena). One hundred ng of DNA was subjected to qPCR to quantify MCMV genome copies (primers ACTAGATGAGCGTGCCGCAT and TCCCCAGGCAATGAACAATC) and human β-actin gene copies (primers GCTGAGGCCCAGTTCTAAAT and TTCAAGTCCCATCCCAGAAAG).

### Antibodies

The following antibodies were used: monoclonal antibodies against GAPDH (14C10; Cell Signaling), β-actin (AC-74; Sigma), HA (3F10; Roche), Flag (M2, Sigma-Aldrich), Cyclin B1 (GNS-1; Santa Cruz), Cyclin E (HE12, Santa Cruz), E2F1 (KH95, Santa Cruz). Antibodies against MCMV IE1 (CROMA101), and MCMV E1 (CROMA103) were provided by Stipan Jonjic (University of Rijeka, Croatia). Polyclonal rabbit antibodies against FAM111B (HPA038637, Sigma-Aldrich), FAM111A (EPR14407, Abcam), Flag (F7425, Sigma-Aldrich), E2F3 (C-18, Santa Cruz) and E2F4 (C-20, Santa Cruz) were used. Secondary antibodies coupled to horseradish peroxidase (HRP) were purchased from DakoCytomation or Jackson ImmunoResearch. Secondary antibodies coupled to Alexa-555 were from Thermo Fischer Scientific.

### Immunofluorescence

10.1 or RPE-1 cells were seeded onto μ-slides (8-well, Ibidi). On the following day, cells were infected with the indicated virus at an MOI of 1 TCID_50_/cell. Cells were fixed with 4% paraformaldehyde in PBS for 20 min at RT, washed with PBS, and free aldehyde groups were blocked with PBS containing 50 mM ammonium chloride for 10 min. Cells were permeabilized by incubation in PBS with 0.3% Triton X-100 for 10 min. PBS containing 0.2% gelatin from porcine skin (Sigma) was used for blocking and dilution of antibodies. Incubations with the antibodies were carried out for 1 h at room temperature (RT). Nuclei were stained for 10 min with DAPI (Roche) or Hoechst 33342 (Invitrogen) at RT. Fluorescence images were acquired by using a Nikon A1+ confocal laser scanning microscope and analysed using ImageJ software.

### Immunoprecipitation and immunoblot analysis

Cells grown in 6-well dishes were infected at an MOI of 2 TCID_50_/cell for 24 h. Lysis of the cells, immunoprecipitation, and immunoblot were performed as previously described (8). Briefly, cells were lysed in a NP-40 buffer (50 mM Tris, 150 mM NaCl, 1% Nonidet P-40, and Complete Mini protease inhibitor cocktail (Roche), Flag-tagged M117 protein was precipitated with anti-Flag and protein G sepharose. Precipitates were washed 3 times with buffer 1 (1 mM Tris pH 7.6, 150 mM NaCl, 2 mM EDTA, 0.2% NP-40), twice with buffer 2 (1 mM Tris pH 7.6, 500 mM NaCl, 2mM EDTA, 0.2% NP-40), and once with buffer 3 (10mM Tris pH 7.6). Precipitated were eluted by boiling in SDS-PAGE sample buffer (125 mM Tris pH 6.8, 4% SDS, 20% glycerol, 10% β-mercaptoethanol, 0.002% bromophenol blue).

For immunoblot analysis, cells were lysed in SDS-PAGE sample buffer. Equal volumes of samples were separated by SDS-PAGE followed by semi-dry transfer to a nitrocellulose membrane (Amersham). Proteins of interest were detected with protein-specific primary antibodies and HRP-coupled secondary antibodies by enhanced chemiluminescence (Amersham) supplemented with 10% Lumigen TMA-6 (Bioquote Limited). All immunoblots shown are representatives of 3 or more experiments.

### Cell cycle experiment

For synchronizing the cells in the G1 phase, 1.7 x 10^4^ RPE-1 cells/cm^2^ were serum starved for 24 h in 0.1% FCS DMEM. To obtain cells in S phase, cells were first synchronized in G1 and then incubated for 24 h with DMEM/10 % FCS containing 1 mM hydroxyurea (Sigma). For G2/M phase arrest, cells synchronized in G1 were incubated for 24 h with DMEM/10% FCS DMEM containing 200 ng/ml nocodazole (Sigma). To synchronize RPE-1 cells in G0, 5.6 x 10^4^ cells/cm^2^ were contact-inhibited and serum starved for 48 h. Asynchronous control cells were grown for 24 h in DMEM/10% FCS. After trypsinization, cells were fixed with 70% (v/v) ice-cold ethanol and kept at 4°C overnight before propidium iodide staining (30 μg/ml propidium iodide + 0.2 mg/ml RNase A in PBS containing 0.1% Triton X-100). Cellular DNA content was measured on a FACSCanto I (Beckton Dickinson) flow cytometer. BD FACS Diva (BD Biosciences) was used to acquire flow cytometric data, and FlowJo (Treestar) was used for analysis.

### RNA sequencing

RPE-1 cells were synchronized for 72 h by serum starvation (DMEM/ 0.5% FCS) and then infected with MCMV at an MOI of 5 in the presence of 10% FCS. Cells were washed 1 hpi to remove input virus and new medium was added. One day post-infection, cells were harvested in RNA lysis buffer and total RNA was extracted using an innuPREP RNA Mini Kit and contaminating DNA was removed as described above. RNA quality was controlled using a Eukaryote Total RNA Nano assay on an Agilent Technologies 2100 Bioanalyzer System. Only high-quality RNA (RIN values between 9-10) was used. Poly(A) mRNA was enriched using the NEBNext Poly(A)mRNA Magnetic Isolation Module (NEB), and cDNA libraries were prepared with a NEXTflex Rapid Directional RNA-Seq Kit (Bioo Scientific) following the manufacturer’s instructions, yielding in library with average cDNA sizes of 500 bp. Libraries were then sequenced on an Illumina HiSeq 2500 system and single-read 50 bp data (single indexed) were generated.

### RNA-seq data analysis

Gene abundance was quantified using salmon software (v1.10.1) (43) with human gene annotations from gencode (version 43) for GRCh38 genome assembly, and imported using R package tximport (v1.28.0) (44). Counts normalization and differential expression (DE) analysis was performed using DESeq2 package (v1.40.2) (45). Null variance of Wald test statistic output by DESeq2 was re-estimated using R package fdrtool (v1.2.17) (46) to calculate *p*-values (and adjusted using the Benjamini-Hochburg method) for the final list of differentially expressed gene. False Discovery Rate (FDR) (BH-adjusted *p*-values) < 0.05 and log2 fold change < ±0.4 was used a criteria for the final DE gene list. Gene ontology enrichment analysis was performed on differentially expressed gene lists using topGO (v2.52.0) (47) GO plots for enrichment of biological process were plotted using R package clusterProfiler (v4.8.3) (48). R statistical computing language (version 4.3.3) was used for the calculations mentioned above except gene level quantification.

### Statistical analysis

Statistical analyses were with the GraphPad Prism 5.0 (Dotmatics) software. Two-way ANOVA with Bonferroni post-hoc test was used for the analysis of qPCR data.

## Acknowledgments

We thank Arne Düsedau (LIV Flow Cytometry Facility) and the NGS facility for technical support. The Leibniz Institute of Virology is supported by the Free and Hanseatic City of Hamburg and the Federal Ministry of Health.

## Data Availability

All data supporting the findings of this study are available within the article.

